# High-resolution wide-field human brain tumor margin detection and in vivo murine neuroimaging

**DOI:** 10.1101/252080

**Authors:** Derek Yecies, Orly Liba, Elliott SoRelle, Rebecca Dutta, Edwin Yuan, Hannes Vogel, Gerald A. Grant, Adam de la Zerda

**Affiliations:** Stanford University Department of Structural Biology; Stanford University Department of Neurosurgery; Stanford University Department of Electrical Engineering; Molecular Imaging Program at Stanford; Bio-X Program at Stanford; Biophysics Program at Stanford University; Applied Physics Program at Stanford University; Stanford Pathology; The Chan Zuckerberg Biohub, San Francisco, CA 94158

## Abstract

Current *in vivo* neuroimaging techniques provide limited field of view or spatial resolution and often require exogenous contrast. These limitations prohibit detailed structural imaging across wide fields of view and hinder intraoperative tumor margin detection. Here we present a novel neuroimaging technique, speckle-modulating optical coherence tomography (SM-OCT), which allows us to image the brains of live mice and *ex vivo* human samples with unprecedented resolution and wide field of view using only endogenous contrast. The increased effective resolution provided by speckle elimination reveals white matter fascicles and cortical layer architecture in the brains of live mice. To our knowledge, the data reported herein represents the highest resolution imaging of murine white matter structure achieved *in vivo* across a wide field of view of several millimeters. When applied to an orthotopic murine glioblastoma xenograft model, SM-OCT readily identifies brain tumor margins with near single-cell resolution. SM-OCT of *ex vivo* human temporal lobe tissue reveals fine structures including cortical layers and myelinated axons. Finally, when applied to an *ex vivo* sample of a low-grade glioma resection margin, SM-OCT is able to resolve the brain tumor margin. Based on these findings, SM-OCT represents a novel approach for intraoperative tumor margin detection and *in vivo* neuroimaging.

## Introduction

Brain tumors are the most common solid tumors in children and the leading cause of pediatric cancer mortality^1^. Extent of resection has been correlated with improved outcome in many types of pediatric and adult brain tumors^2–11^. However, maximal safe resection is often limited by an inability to distinguish tumor from normal brain. To address this problem, a variety of intraoperative imaging modalities have been investigated, including magnetic resonance imaging (MRI)^12–15^, wide-field fluorescence^7^, high-resolution fluorescence microscopy (confocal and multiphoton imaging)^16^, and optical coherence tomography (OCT)^17–21^.

Intraoperative MRI (iMRI) allows for the reliable identification of macroscopic residual tumor and has been shown to increase the extent of resection in glioma surgery^14,15,22,23^, however, iMRI has several limitations. iMRI systems are expensive and require the use of specialized non-ferromagnetic surgical equipment. The spatial resolution of MRI precludes the identification of small regions of residual tumor. Furthermore, the acquisition of iMRI is time consuming and requires a complete cessation of surgery for image acquisition.

Wide-field fluorescence guided brain tumor surgery, most notably with 5-aminolevulinic acid (5-ALA), has been shown to improve extent of resection for patients with glioblastoma and is easily incorporated into a standard surgical workflow^7^. However, wide-field fluorescence imaging relies on the presence of a disrupted blood brain barrier, and in the case of 5-ALA, specific biochemical alterations that are not present in all tumors^24,25^. Therefore, the utility of wide-field fluorescence guided brain tumor surgery outside of glioblastoma has been limited, especially in low grade glial tumors and pediatric tumors that may have an intact blood brain barrier^26–29^. Confocal microscopy allows for the imaging of 5-ALA fluorescence in some low-grade gliomas^16^. Additionally, label-free confocal and multiphoton microscopy systems are able to resolve individual axons and *ex vivo* brain tumor margins^30–34^. However the requirement for tissue contact, limited field of view, and limited depth of penetration inherent to confocal and multiphoton microscopy may limit their practical utility^35^.

OCT allows for rapid, wide-field, and label-free *in vivo* brain imaging with microscopic resolution and up to two millimeters of tissue penetration^36–38^. Additional features that make OCT particularly appealing for intraoperative use include the absence of photobleaching seen with fluorescence-based modalities, a safe low-energy infra-red light source, and the ability to integrate with the existing optics of a standard neurosurgical operating microscope. Several studies have demonstrated the feasibility of building OCT systems into neurosurgical operating microscopes and endoscopes and the ability to successfully differentiate tumor from normal brain based on signal attenuation and the presence of gross structural features within the tumor^19,21,39–41^.Further work has established OCT attenuation thresholds that may allow for the rapid differentiation of tumor from cerebral white matter, though attenuation alone was not able to differentiate tumors from grey matter^42,43^. While promising, the effective resolution and image quality in these studies were significantly degraded by the presence of speckle noise that is intrinsic to OCT imaging.

*in vivo* neuroimaging studies with optical coherence microscopy (OCM), a closely related modality that utilizes high-magnification objective lenses in conjunction with OCT, are suggestive of the image quality that can be obtained when speckle noise is eliminated from OCT imaging. The high magnification of OCM systems allows for the utilization of spatial averaging to eliminate speckle noise while still maintaining micrometer-scale resolution. After the removal of speckle, OCM is capable of *in vivo* imaging of individual axons and pyramidal cell bodies as well as the laminated architecture of cortical layers in mice and rats^44–46^. However, the narrow field of view, need for contact of the imaging objective with the sample, and shallow depth of field, requiring complex scanning of the image focal plane and extremely stable imaging samples, limit the potential of OCM for intraoperative use.

In this paper, we introduce a novel method for label-free *in* vivo neuroimaging and tumor margin detection, Speckle-Modulating OCT (SM-OCT).SM-OCT allows for the significant reduction of speckle noise within OCT images without compromising spatial resolution, effectively increasing the resolution of standard OCT imaging while maintaining imaging speed, depth of field, and field of view^47^. In this manuscript, we demonstrate that SM-OCT is capable of high-resolution structural imaging of the mouse brain, which is unprecedented for wide-field *in vivo* imaging. Specifically, individual white matter fascicles of the cingulum bundle, corpus callosum and alveolus of the hippocampus are clearly visible across a wide field of view using only the endogenous contrast of neural tissue. Fine structures of the brain including myelinated axons and cortical layers are also clearly visualized. *in vivo* SM-OCT imaging of mice with orthotopic glioblastoma xenografts identifies tumor margins with near cellular level specificity. Finally, when applied to *ex vivo* human samples, SM-OCT allows for label-free delineation of the brain tumor margin of a low-grade glioma. Our work suggests that fast, wide field of view, high-resolution *in vivo* optical imaging using SM-OCT has the potential to significantly advance the care of neurosurgical patients and expand the capabilities of small animal neuroimaging.

## Results

### SM-OCT *in vivo* Neuroimaging

Anesthetized BL6 mice with glass cranial windows underwent wide-field brain imaging with OCT and SMOCT (Fig. 1 and 2). OCT imaging penetrates brain tissue to a depth of 1.5mm and can resolve the gross structure of the hippocampus and corpus callosum, however speckle noise precludes the resolution of further structural detail (Fig. 1b, 2a, 2d, and S2). SM-OCT significantly reduces speckle noise, revealing the fine anatomical structure of the brain including cortical layers (Fig. 1) and white matter fascicles (Figure. 2).

Mammalian cortex has four to six distinct layers that are differentiated both histologically and functionally. SM-OCT allows for clear visualization of the architecture of the cortical layers (Fig. 1c). The underlying signal characteristics of mouse brain imaged with SM-OCT demonstrate significant noise reduction compared to the OCT image, with distinct variability between the signal intensity among the different cortical layers allowing for their visualization (Supplementary Fig. 1). The signal differences between layers are consistent with prior literature from OCM and are explained by differences in the density of cells and neuritic processes between layers, with more cell-body dense layers such as layer two being darker and more axon dense layers such as layer one being brighter^46,48,49^.

**Figure 1:**
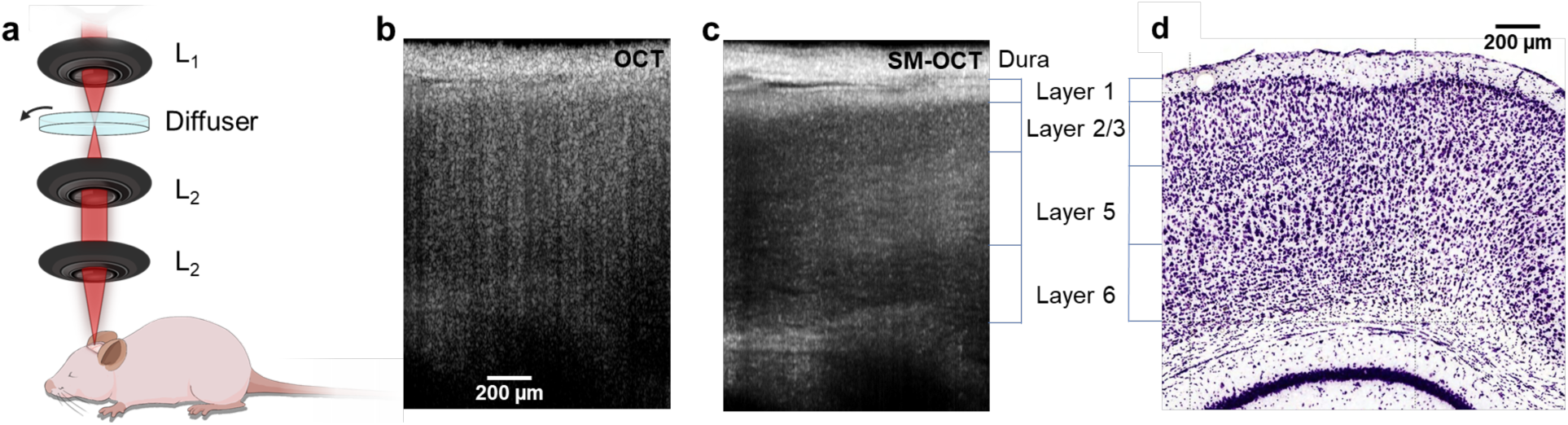
SM-OCT imaging of mouse brain *in vivo* reveals cortical layers. a) The sample arm the SM-OCT system, L_1_ is the main lens of the OCT, the diffuser is rotated in the focal plane, which is relayed by two lenses, L_2_, in a 4f configuration. b) OCT B-scan of mouse cortex. c) SM-OCT B-scan of mouse cortex, showing the cortical layers, which are revealed by removing the speckle noise. d) Histology of mouse brain (image credit: Allen Institute ^56^), showing the corresponding cortical layers to SM-OCT.

**Figure 2:**
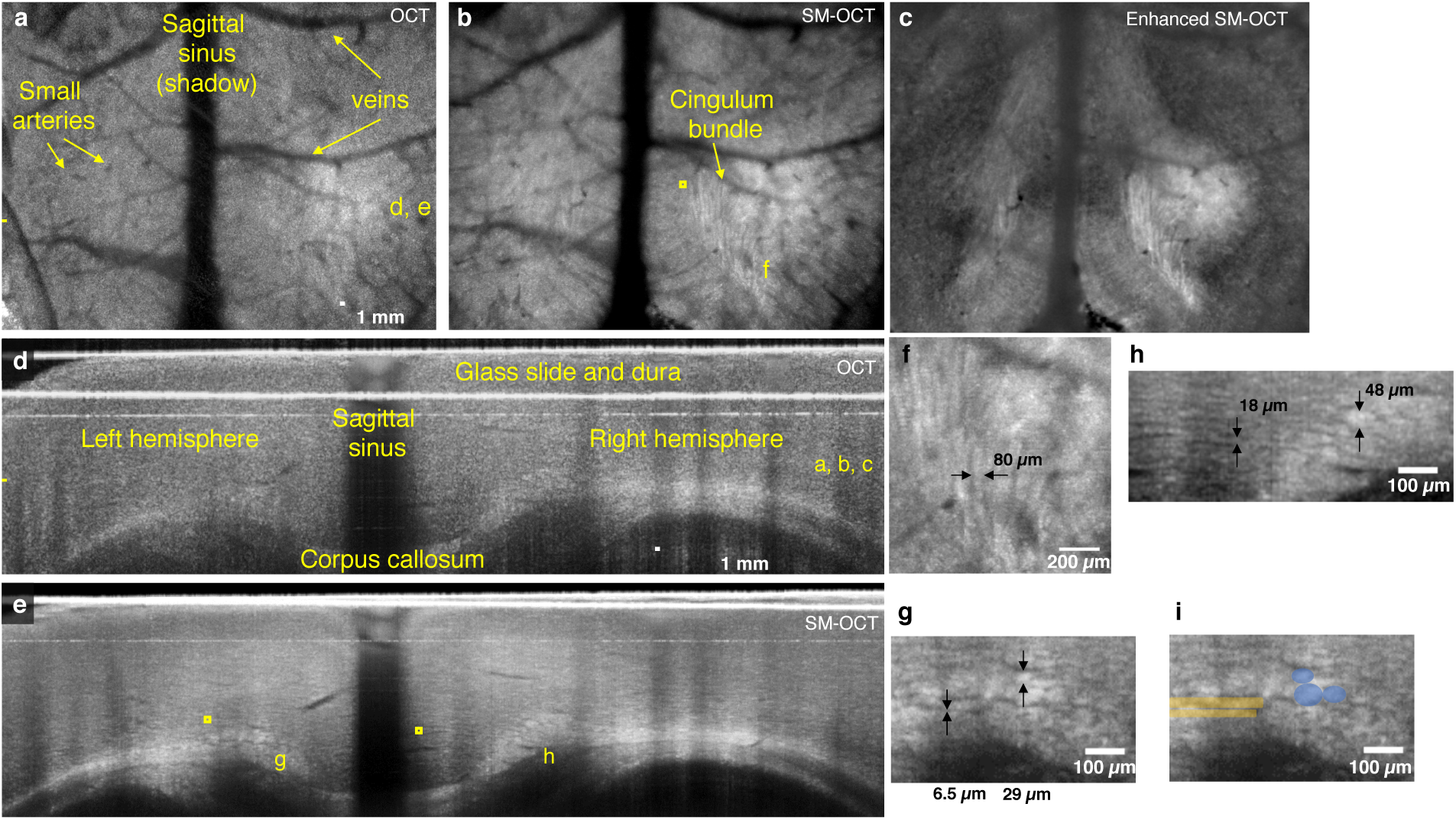
SM-OCT reveals white matter fascicles in mice *in vivo*. a) OCT axial view of mouse cortex, depth is shown as yellow dashed line in (d). b) SM-OCT image of the region showed in (a). The removal of speckle reveals white matter structures, including the cingulum bundle. c) The white matter structures shown by SM-OCT can be enhanced by image processing. d) OCT coronal view of mouse cortex, the location is shown as yellow dashed line in (a). SM-OCT image of region shown in (d), revealing the white matter structures in high-resolution. f) A close-up axial view of the cingulum bundle. g, h) close-up coronal views of white matter structures of various sizes including the tracts of the cingulum bundle and very small unnamed fascicles. i) close-up coronal view with manual segmentation of several fibers of cingulum bundle (purple) and very small unnamed fascicle (yellow).

Small individual white matter fascicles of the cingulum bundle, corpus callosum, and alveolus of the hippocampus are clearly visualized *in vivo* for the first time with optical imaging using SM-OCT (Fig. 2b, 2e-h, Supplementary Fig. 2b, 2d, Movie 1-3. These fascicles are approximately 40-80 um in diameter (Fig 2f-g), and cannot be resolved using OCT (Supplementary Fig 2a, 2c, Movie 3). To our knowledge, these structures have not been directly visualized *in situ* in live animals prior to this study. Three-dimensional imaging of these structures allows for them to be visualized in multiple planes, for example individual fascicles of the cingulum bundle are seen as long tubes in the axial plane (Fig. 2b and 2f, Supplementary Fig. 2b), and as circles in the coronal cross section (Fig. 2e, 2g-h and Supplementary Fig. S2d). These structures may also be manually segmented (Fig 2i).

Closer examination of coronal plane SM-OCT images reveals even smaller structures, whose size (6-18 um) and location are consistent with individual myelinated axons or very small fascicles consisting of a small number of myelinated axons (Fig. 2h). These structures can also be seen coursing from the medial cortex into the cingulum bundle in the axial plane (Supplementary Movie 2).These structures cannot be seen with OCT due to speckle noise (Supplementary Fig. 2c and Supplementary Movie 3). As a wide-field *in vivo* neuroimaging platform, SM-OCT can resolve a wide range of tissue features from cortical layers to small white matter fascicles up to 1.5 mm deep within tissue using only endogenous contrast.

Unenhanced SM-OCT neuroimaging allows for exquisite *in vivo* imaging of the murine brain. However, SM-OCT data also lends itself to digital enhancement. To achieve this, we applied vessel shadow removal, anisotropic diffusion denoising^50^, and contrast enhancement to the SM-OCT volumes (Fig. 2c and Supplementary Movie 4). This enhancement allows improved visualization of the axonal and white matter architecture by enhancing edges and contrast.

### High-resolution *in vivo* imaging of orthotopic murine xenograft brain tumors

We next applied SM-OCT to an orthotopic glioblastoma model. Nude mice were implanted with U87 glioblastoma cells and prepared with glass cranial windows. The mice were imaged with SM-OCT on post-implantation day seven (Fig. 3 and Supplementary Movie 5-7). The wide field of view achievable with SMOCT allows for the entire tumor and surrounding brain tissue to be imaged. SM-OCT clearly differentiates tumor from normal brain based on signal intensity (Fig. 3). The undisturbed regions of the brain are rich in highly reflective myelin and appear bright, while the tumor appears dark because it lacks myelinated structures and includes mainly cell bodies that are significantly less reflective^46,48,49^.The high resolution of SM-OCT allows for imaging of the brain tumor margin with near single cell resolution, including visualization of the finger-like projections of the tumor growing along axonal projections (Fig. 3e). In the OCT images (Supplementary Fig. 3), the tumor margin is less visible due to speckle noise.

**Figure 3:**
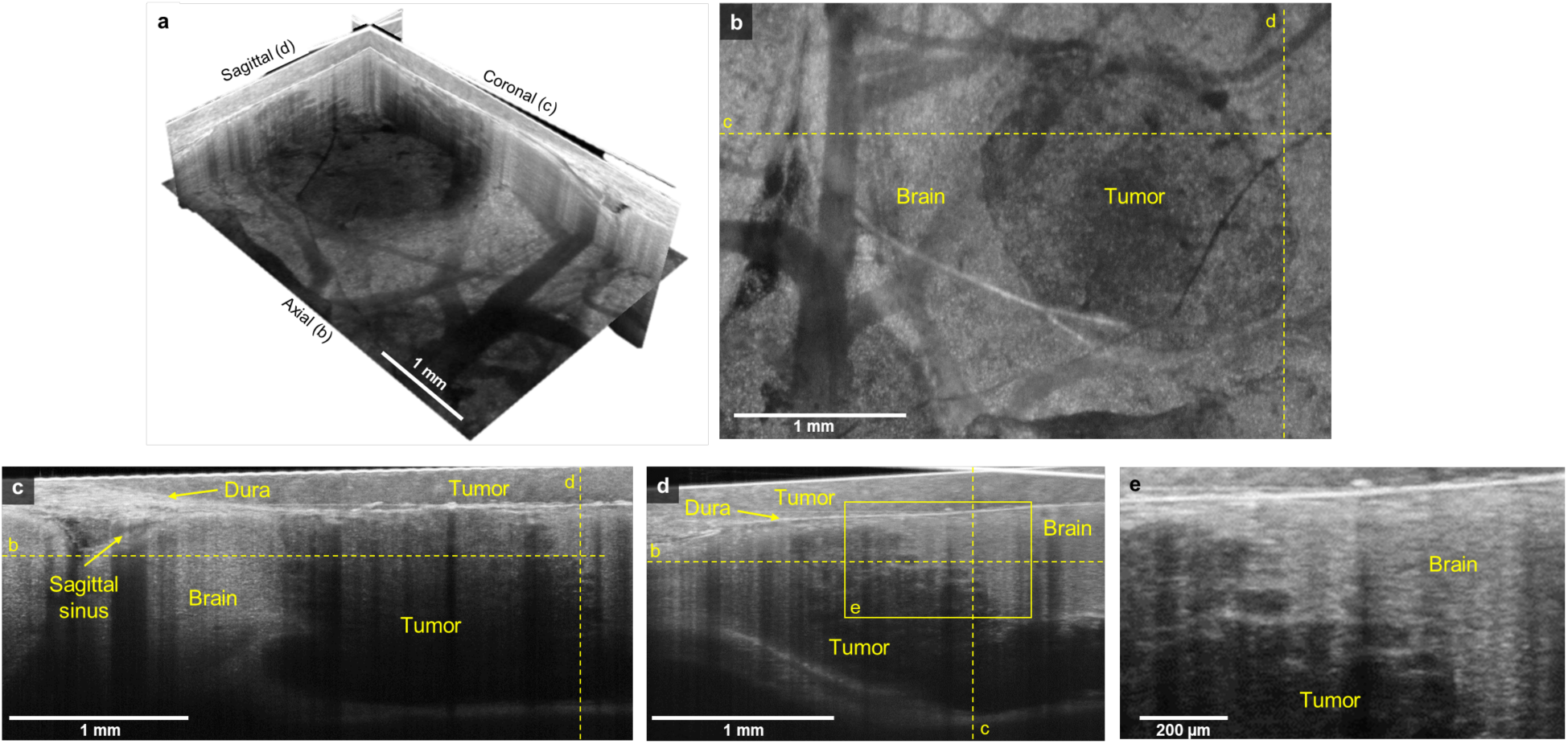
SM-OCT reveals high-resolution features of tumor margin *in vivo*. a) SM-OCT ortho-slice of the tumor volume, showing the different sections in three dimensions. b) SM-OCT axial view of mouse cortex with a GBM tumor, depth is shown as yellow dashed line in (c, d). c, d) SM-OCT coronal and sagittal views, respectively, showing the tumor margin, the locations are shown as yellow dashed lines in (b). e) A close-up view of the tumor margin in (d), showing the finger-like invasion of the into the surrounding brain tissue.

### Imaging of human brain *ex vivo*

Given the promising results of SM-OCT in murine neuroimaging, we investigated whether similar results could be obtained in human brain tissue. A fresh, unfixed section of the inferior temporal gyrus from a temporal lobectomy was taken directly from the operating room to our laboratory for imaging with OCT and SM-OCT. Similar to *in vivo* murine imaging, OCT was able to image human brain tissue to a depth of greater than 1.5mm. However, speckle noise in the OCT image prevents the resolution of cortical layers or axonal processes (Fig. 4a). SM-OCT clearly resolves the first three cortical layers as well as individual myelinated axonal processes, primarily in layers one and three (Fig. 4b, 4d, and 4e). Axons exist in all cortical layers. However, since SM-OCT is dependent on reflected light, axons oriented perpendicular to the incident beam, such as those in layers one and three provide significantly greater signal, allowing them to be resolved with this method.

**Figure 4:**
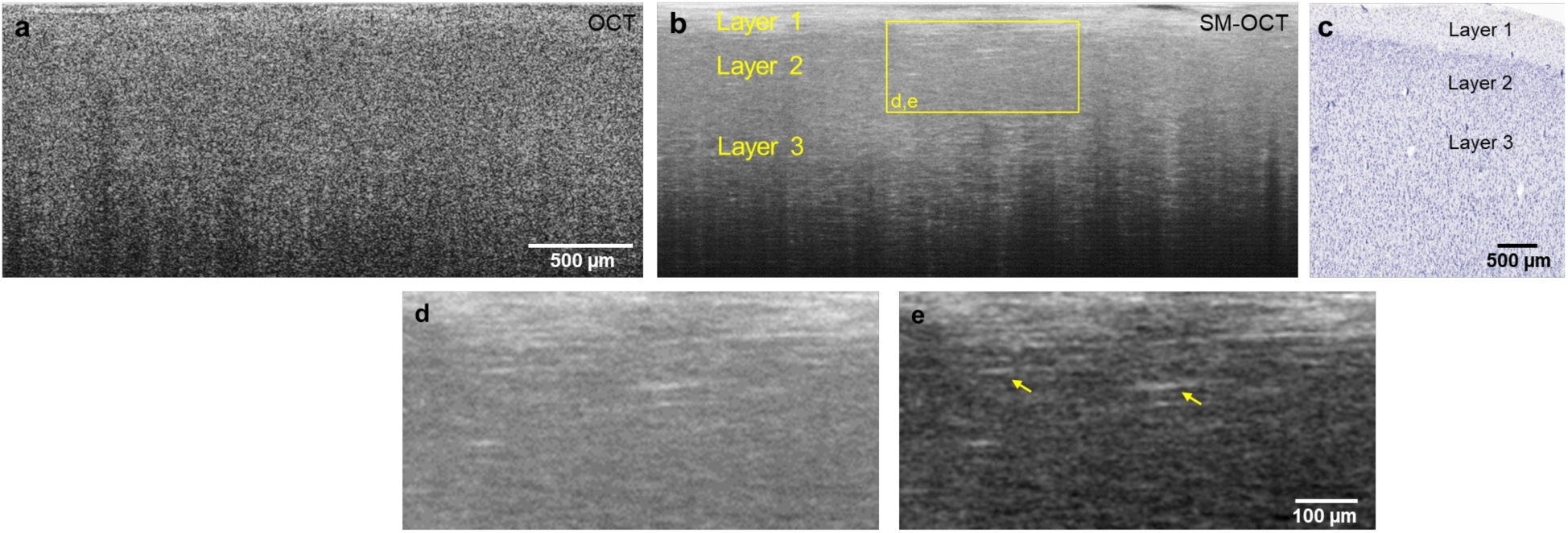
SM-OCT of *ex vivo* human brain sample reveals cortical layers and axons. a, b) OCT and SM-OCT B-scans of cortex. The SM-OCT image reveals cortical layers and myelinated axonal projections. c) Corresponding histology (image credit: Allen Institute ^57^). d, e) a close-up view of the myelinated axons shown in b. The contrast in e is enhanced to highlight the myelinated axons.

### Imaging of human low-grade glioma margins *ex vivo*

Encouraged by the capabilities demonstrated by SM-OCT with *in vivo* murine and *ex vivo* human brain imaging, we sought to apply SM-OCT to the substantial clinical problem of tumor margin detection in low-grade gliomas. A child with a low-grade glioma and epileptic foci extending beyond the tumor underwent a planned supratotal resection of the tumor and adjacent epileptic regions. A fresh, unfixed sample that contained the presumed tumor margin based on stereotactic image guidance was taken directly from the operating room to our laboratory for imaging with SM-OCT and was subsequently fixed for histological processing. Alignment of the sample was preserved throughout imaging and histologic processing. Histologic sections were aligned with b-scans from the three-dimensional SM-OCT volume based on vessel anatomy (Fig. 5 a-c). The tumor tissue in the SM-OCT image demonstrates lower intensity signal compared to the brain tissue as well as a loss of normal structures, such as layer one axons, seen in the normal brain (Fig. 5a and 5b). The subtle tumor margin seen in the histologic sample (Fig. 5a) is recapitulated in the SM-OCT image (Fig. 5b) and rendered quite clear with by stretching the brightness and contrast of the SMOCT image (Fig. 5c). When the same sample is visualized from an axial/top down plane, following the flattening of the volume, several important features are apparent. Individual myelinated axonal processes in layer one of the healthy brain are clearly visualized on the left (Fig. 5d-f), while on the right side of the image, layer one axons are absent and have been replaced by the tumor (Fig 5d and 5f). Thus, the presence or absence of layer one axons provides an additional means by which SM-OCT can resolve the tumor margin of this low-grade glioma with high spatial resolution across a field of view of several millimeters.

**Figure 5:**
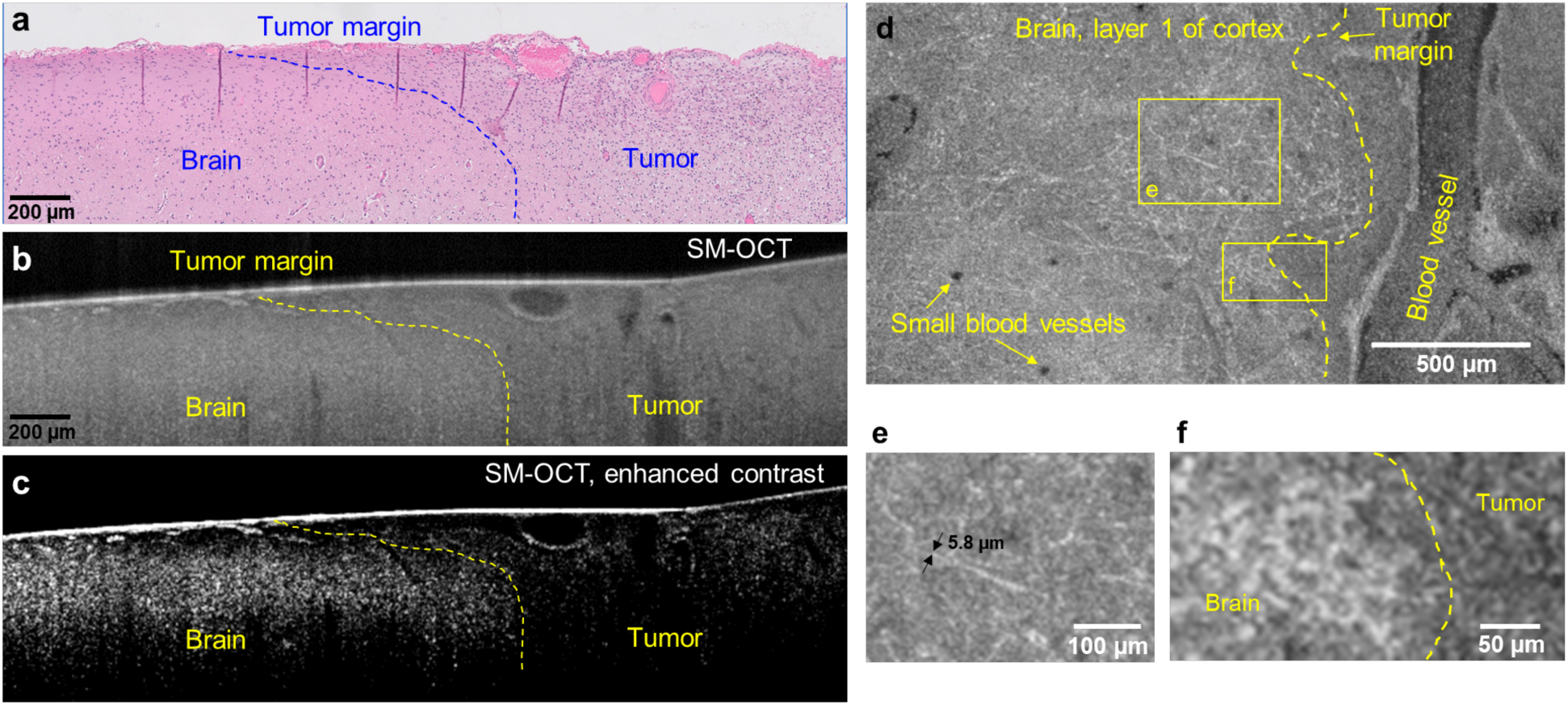
*ex vivo* human LGG tumor margin visualized with histology and SM-OCT. a) Histology shows the difference in cell density and arrangement in the brain (sparse and organized) and tumor (dense and disorganized) regions, the tumor margin can be roughly estimated and is shown by the blue dashed line. b) SMOCT B-scan of the brain sample, corresponding to (a), showing the tumor as having reduced signal intensity compared to the normal brain, likely due to the reduced optical scattering of the non-myelinated tumor cells. c) The image in (b) after contrast-enhancement, emphasizing the difference in signal intensities between the tumor and brain regions. d) SM-OCT axial view (*en face*) of the tumor margin, demonstrating invasion of layer 1 of the cortex by the glioma. The image shows the axonal projections of layer 1 at the left of the tumor margin and the lack of them at the region of the tumor. e) A close-up view on the axonal projections of layer 1 of cortex. The thickness of the measured myelinated axons is limited by the optical resolution of the OCT. f) A close-up view on the tumor margin, highlighted by the lack of axons and lower signal intensity in the tumor.

## Discussion

In this proof-of concept paper we demonstrate that SM-OCT can address fundamental shortcomings of current *in vivo* neuroimaging modalities used for small animal neuroscience and intraoperative tumor margin detection. Specifically, SM-OCT can resolve brain tumor margins and neural structures ranging from individual axons to white matter fascicles and cortical layers with a resolution of a few microns across a wide field of view of several millimeters without the use of exogenous contrast agents. To our knowledge, there are no other imaging modalities that currently occupy this crucial space in *in vivo* imaging, and accordingly SM-OCT neuroimaging lends itself to a myriad of scientific and clinical uses.

SM-OCT could allow for serial *in vivo* imaging in murine models of neurologic disease such as alzheimer’s disease, epilepsy, and stroke. A major advantage of serial imaging is that it allows for the disease process within a single mouse to be tracked over time, in response to treatments, and across different stages of disease. The resolution provided by SM-OCT may yield new insights into neurologic disease processes, as current murine studies that utilize serial *in vivo* imaging rely on MRI or PET, which have inherent resolution limitations. The design of these studies would need to be mindful of the limitations of SM-OCT, primarily the limited depth of penetration of 1-1.5mm and a relative difficulty in imaging anisotropic structures oriented parallel to the incident beam due to their reduced reflectivity.

The ability of SM-OCT to resolve brain tumor margins with near single-cell resolution has exciting applications for intraoperative use. There are currently no intraoperative imaging modalities capable of differentiating certain tumor types, such as low-grade gliomas, from normal brain, and the ability of SMOCT to accomplish this without the use of exogenous contrast is likely to be of significant value to neurosurgeons and patients with neurosurgical diseases. The SM-OCT setup described in this paper is simple, robust, and can likely be integrated with the optics of a standard operating microscope with relatively minor modifications. Alternatively, SM-OCT could be used on *ex vivo* samples taken during an operation, similarly to what was done in this paper, to provide information comparable to a frozen section, the current gold standard for intraoperative tissue diagnosis.

OCT technology continues to develop rapidly and future iterations of SM-OCT will likely be able to take advantage of advances made across the field. For example, the SM-OCT system used in this paper can scan up to a relatively fast rate of 91khz. However, there are currently swept source OCT systems that are capable of scanning at rates more than an order of magnitude faster^51^. These systems could achieve volumetric video-rate imaging and have the potential to enable surgeries that are done under continuous OCT visualization^52^. Additionally, swept source systems often utilize higher powered light sources that could increase the imaging depth in tissue^37^. Polarization sensitive OCT has been utilized in *ex vivo* analysis of thin slices of brain tissue and provides high-resolution visualization of white matter structures owing to their variable directionality^53–55^. The modification of SM-OCT for swept source and polarization sensitive OCT systems and other varieties of OCT beyond simple spectral domain OCT represents an interesting future area of investigation.

In future experiments, we plan to utilize SM-OCT in the operating room to further investigate the role of this technology for intraoperative use. The ability to resolve cortical layer architecture and patterns of axonal arborization may allow for uses beyond tumor margin detection, such as identification of areas of cortical dysplasia to improve the accuracy of epilepsy surgery. The human tissue experiments presented in this paper attempted to mirror intraoperative conditions as much as possible by utilizing fresh, unfixed tissues taken immediately from the operating room to the lab for imaging. As such, we are hopeful that the results presented in this paper will be reproduced in future intraoperative experiments.

## Methods

### Cranial window preparation and orthotopic brain tumor animal models

The animal experiments were carried out in accordance with Stanford University Institutional Animal Care and Use Committee guidelines under APLAC protocols 26548 and 27499. Female athymic nude mice (*Foxn1^nu-/nu-^*, Charles River Labs) were anesthetized with ketamine/xylazine (100/10 mg/kg, intramuscular injection) and placed in a stereotactic frame. The scalp and underlying soft tissue over the parietal cortex were removed bilaterally. A drill was used to create a rectangular cranial window centered on the midsagittal suture that extended from the bregma to the lambdoid sutures. U87 human xenografts were implanted (10 μL of 1 x 10^6^ cells/ml cell suspension). Following implantation, a 5.5 mm x 7.5 mm x 3 mm glass cranial window was glued to the bone surrounding the cranial window with cyanoacrylate. Female BL6 mice were prepared using the same protocol, but without the injection of tumor cells.

### Imaging system

All OCT images were acquired using a commercial OCT which was adapted to perform speckle-modulation (SM-OCT)^47^ for the removal of speckle noise. The commercial OCT was a spectral-domain system (Telesto, ThorLabs, Newton, NJ), with a center wavelength of 1300 nm and 170 nm bandwidth, which provide and axial resolution of 3.7 μm in tissue. The spectrometer of the OCT acquires 1024 samples for each A-scan at a rate of 28 kHz. Speckle modulation was implemented by inserting a rotating ground glass diffuser at a conjugate image plane. The main lens of the OCT provides a lateral resolution of 9 μm (FWHM) and depth of field (DOF) of 270 μm in water (LSM03, ThorLabs, Newton, NJ). The ground glass diffuser was custom made by lapping a 3 mm thick glass window with AR coating at the wavelength range of the OCT (ThorLabs, Newton, NJ). The window was ground by lapping with 5 μm aluminum oxide grit (Universal Photonics, Central Islip, NY) for 15 minutes. The diffuser was rotated using an electromechanical mount (RSC-103, Pacific Laser Equipment, Santa Ana, CA). The optic axis was aligned with a radial offset relative to the center of the diffuser such that rotation ensured changing speckle patterns in the incident beam. The image plane was replicated by two lenses (LSM02, ThorLabs, Newton, NJ) in a 4F configuration. Traditional OCT volumes were acquired in addition to SM-OCT volumes, by removing the diffuser from the optical axis.

### Imaging protocol

BL6 mice were imaged at least seven days after the implantation of the cranial window. Nude mice were imaged seven days after tumor-cell and window implantation. The mice were anesthetized with 2% isoflurane and placed in a stereotactic frame.

SM-OCT and OCT 3D volumes were acquired with 6 μm lateral spacing and 18 to 40 averages with an A-scan rate of 28 kHz. For SM-OCT acquisition, the diffuser was rotated at a tangential speed of ∼2 mm/s, which created non-correlated speckle patterns. 2D B-scans (figures 1 and 4) were acquired with 4 μm lateral spacing and 100 averages with an A-scan rate of 28 kHz.

### OCT and SM-OCT post-processing

All processing and analysis were performed with Matlab (MathWorks, Natick, MA). Visualization of volumes was done using ImageJ^58^.

The structure of the tissues was obtained by a conventional OCT reconstruction algorithm with custom dispersion compensation^59,60^. SM-OCT provides frames with uncorrelated speckle patterns, which were intensity-averaged in linear-scale. The images and volumes were uniformly re-sampled to obtain a cube-shaped voxel with 4 μm sides.

The volume of mouse brain (Fig. 2c, Supplementary Movie 4) was further post-processed to enhance the visibility of the white-matter fascicles. First, the low signal of the blood vessels and their shadows was increased to the mid-range of the signal-scale. This was done by applying a compensating gain to the axial (*en face*) image so that the average signal along the depth dimension is uniform along the field of view. Next, three-dimensional anisotropic diffusion^50^ was applied to the volume. Last, the contrast was enhanced using histogram equalization.

### Histology images

Histology images of mouse and human brain in figures 1 and 4, respectively, were obtained from the Allen Institute website. The image of mouse cortical layers is found in: http://atlas.brain-map.org/atlas?atlas=1#atlas=1 &structure=451 &resolution=11.79 &x=5536.251100352113 &y=4142.8608 39736294 &zoom=-3 &plate=100960240

The image of human cortical layers is found in:http://atlas.brain-map.org/atlas?atlas=138322605#atlas=138322605 &structure=12142 &resolution=7.77 &x=54524 &y=822 12 &zoom=-3 &plate=112360933 &z=8.

The human tumor margin sample in figure 5 was fixed in formalin, paraffin embedded and stained with hematoxylin and eosin with care taken to preserve the orientation in which the sample was imaged with OCT. The tissue slides were then imaged with a bright-field microscope. All human tissue was obtained following IRB exemption for this study and was performed under the auspices of the existing Stanford Brain Bank IRB protocol.

## Acknowledgments

This work was funded in part by grants from the Claire Giannini Fund, the United States Air Force (FA9550-15-1-0007), the National Institutes of Health (NIH DP50D012179), the National Science Foundation (NSF 1438340), the Damon Runyon Cancer Research Foundation (DFS#06-13), the Susan G. Komen Breast Cancer Foundation (SAB15-00003), the Mary Kay Foundation (017-14), the Donald E. and Delia B. Baxter Foundation, the Skippy Frank Foundation, the Center for Cancer Nanotechnology Excellence and Translation (CCNE-T; NIH-NCI U54CA151459) and the Stanford Bio-X Inter-disciplinary Initiative Program (IIP6-43). A.d.l.Z is a Chan Zuckerberg Biohub investigator and a Pew-Stewart Scholar for Cancer Research supported by The Pew Charitable Trusts and The Alexander and Margaret Stewart Trust. O.L. is grateful for a Stanford Bowes Bio-X Graduate Fellowship.D.Y is greatful for the funding support for this publication provided by the Tashia and John Morgridge Endowed Postdoctoral Fellowship from the Child Health Research Institute at Lucille Packard Children’s Hospital as well as the National Institute of Neurological Disorders and Stroke of the National Institutes of Health under Award Number R25NS065741. The content is solely the responsibility of the authors and does not necessarily represent the official views of the National Institutes of Health. E.D.S. wishes to acknowledge funding from the Stanford Biophysics Program training grant (T32 GM-08294). We thank Christy Wilson PhD for training regarding the preparation of murine cranial windows, Andrew Olson and the Stanford Neuroscience Microscopy Service (supported by NIH NS069375) for access to Imaris x64 software, Timothy R. Brand and the Ginzton Crystal Shop for creating the lapped diffuser and Nick A. Suhar for custom artwork in Fig. 1a.

## Supplementary Figures

**Supplementary Figure 1:**
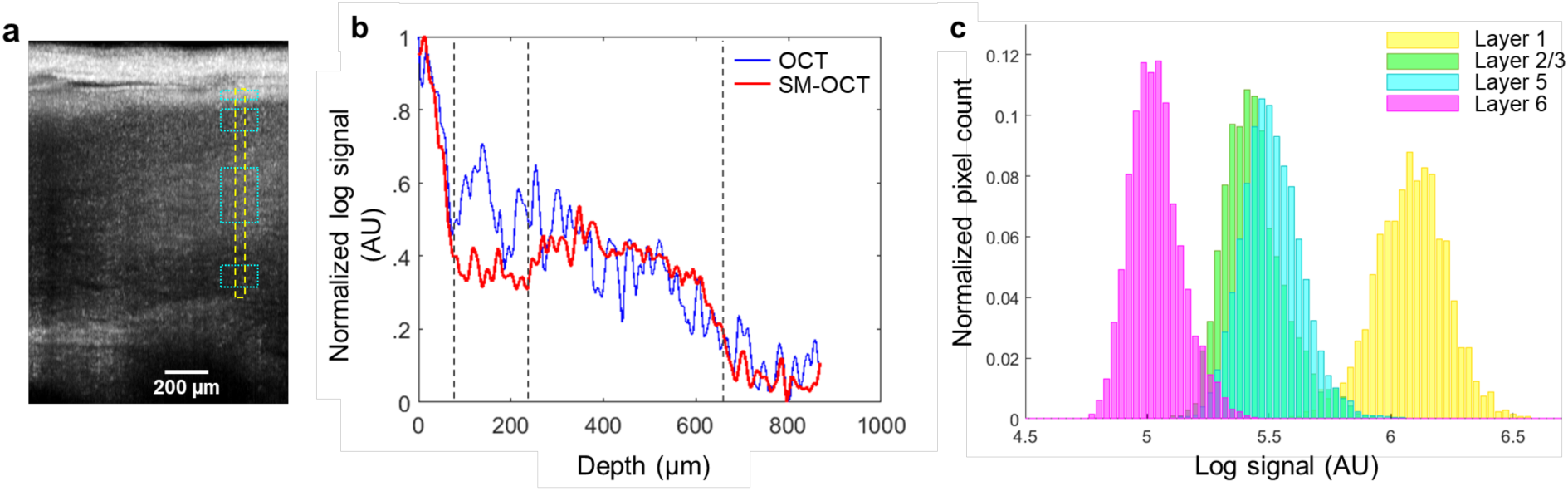
SM-OCT imaging of mouse brain *in vivo* reveals cortical layers. a) SM-OCT B-scan of mouse cortex, showing the areas used for quantifying the signal in the following analysis. b) the signal intensity of OCT and SM-OCT as a function of depth. The region of the signal is shown as a yellow dashed line in (a). c) Normalized histograms of the SM-OCT signal intensity, on the regions shown by cyan rectangles in (a).

**Supplementary Figure 2:**
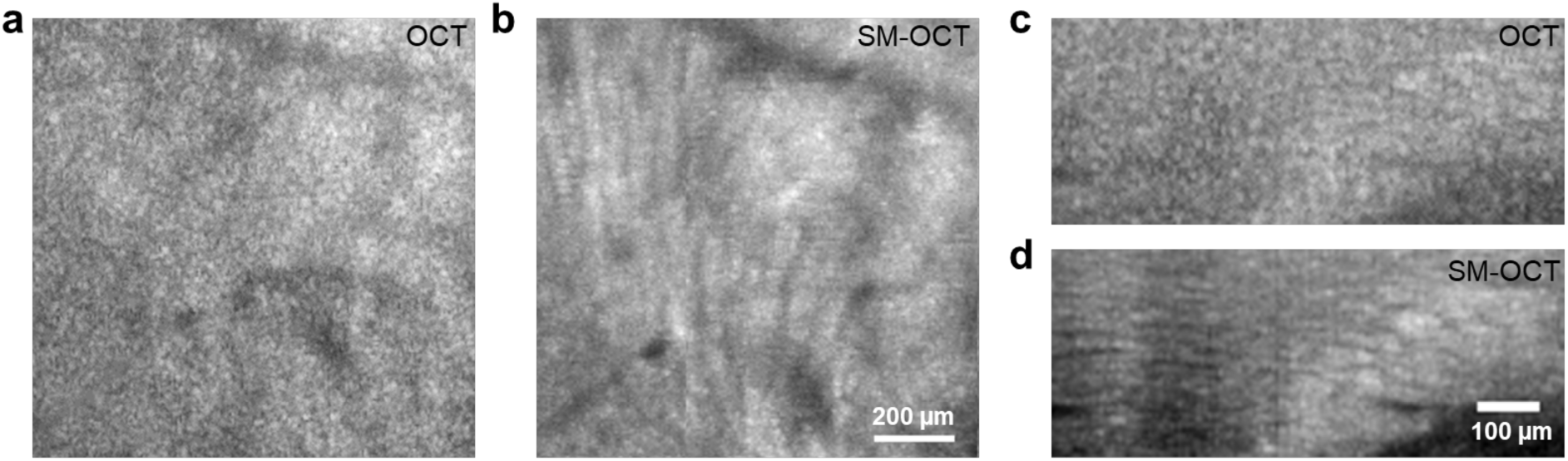
Comparison between SM-OCT and OCT images of mouse white-matter structures *in vivo*. a, b) OCT and SM-OCT (respectively) axial close up views (*en face*) of the cingulum bundle. The structures of the fascicles are revealed through speckle removal. c, d) OCT and SM-OCT (respectively) coronal close up views (B-scans) of the white matter structures, revealed in high-resolution when using SM-OCT.

**Supplementary Figure 3:**
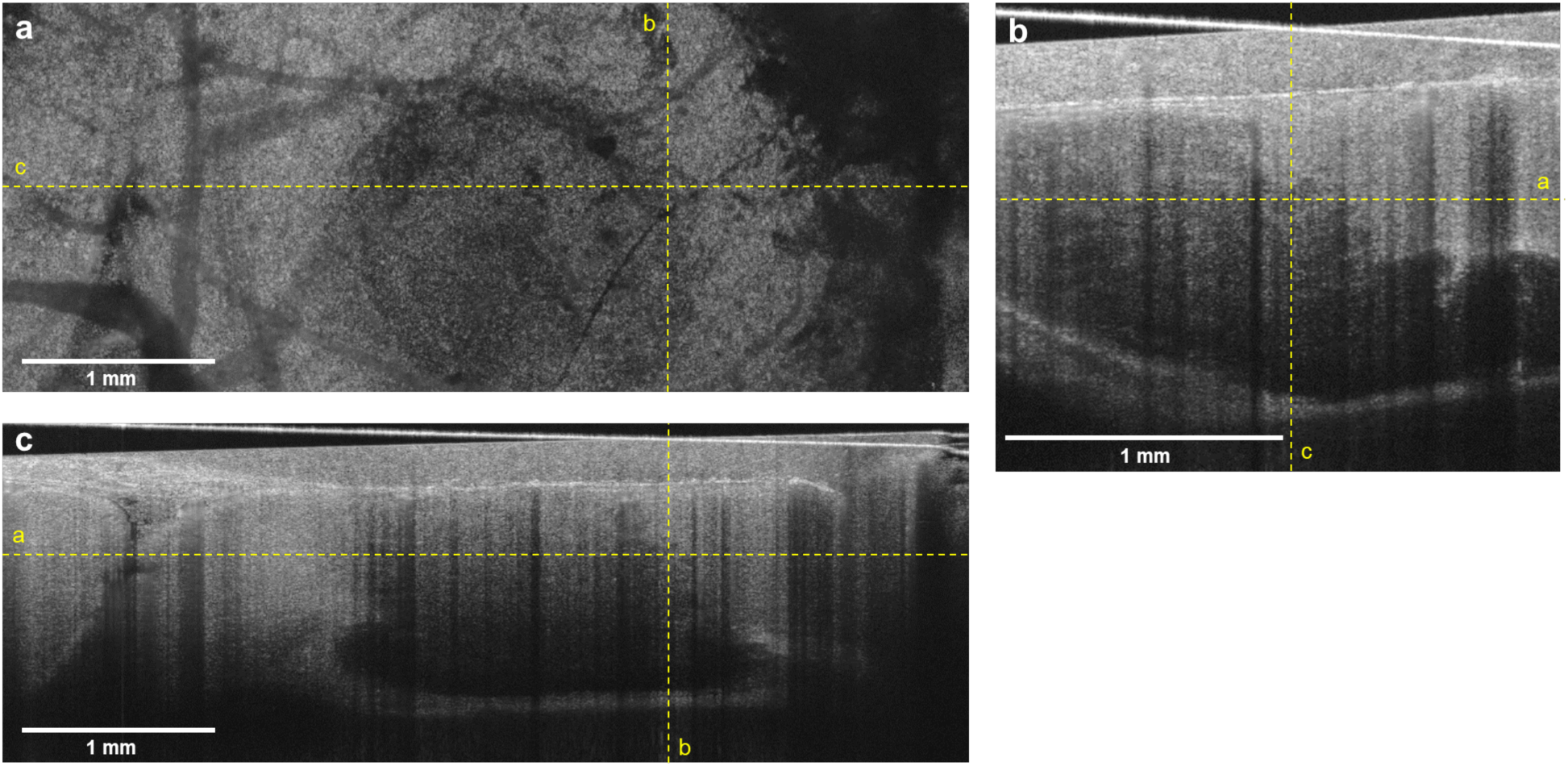
OCT volume, corresponding to the SM-OCT volume in Fig. 3. Axial (a), sagittal (b) and coronal (c) views of the GBM tumor margin, imaged with OCT *in vivo*. Compared to the SM-OCT volume, the speckle noise in OCT obscures the fine structures at the tumor margin.

**Supplementary movie captions**

